# Dose-dependent changes in global brain activity and functional connectivity following exposure to psilocybin: a BOLD MRI study in awake rats

**DOI:** 10.1101/2025.02.01.636078

**Authors:** Evan Fuini, Arnold Chang, Richard J. Ortiz, Taufiq Nasseef, Josh Edwards, Marc Latta, Elias Gonzalez, Taylor J. Woodward, Heather B. Bradshaw, Bryce Axe, Ashwath Maheswari, Noah Cavallaro, Praveen P. Kulkarni, Craig F. Ferris

## Abstract

Psilocybin is a hallucinogen with complex neurobiological and behavioral effects. This is the first study to use MRI to follow functional changes in brain activity in response to different doses of psilocybin in fully awake, drug naive rats. Female and male rats were given IP injections of vehicle or psilocybin in doses of 0.03 mg/kg, 0.3 mg/kg, and 3.0 mg/kg while fully awake during the imaging session. Changes in BOLD signal were recorded over a 20 min window. Data for resting state functional connectivity were collected approximately 35 min post injection All data were registered to rat 3D MRI atlas with 173 brain regions providing site-specific changes in global brain activity and changes in functional connectivity. Treatment with psilocybin resulted in a significant dose-dependent increase in positive BOLD signal. The areas most affected by the acute presentation of psilocybin were the somatosensory cortex, basal ganglia and thalamus. Females were significantly more sensitive to the 0.3 mg/kg dose of psilocybin than males. There was a significant dose-dependent global increase in functional connectivity, highlighted by hyperconnectivity to the cerebellum. Brain areas hypothesized to be involved in loss of sensory filtering and organization of sensory motor stimuli such as the claustrum and the cortico-basal ganglia-thalamic-cortical loop were all affected by psilocybin in a dose-dependent manner. Indeed, the general neuroanatomical circuitry associated with the psychedelic experience was affected but the direction of the BOLD signal and pattern of activity between neural networks was inconsistent with the human literature.

## Introduction

Psilocybin (PSI) has gained interest in psychological research in recent years for its potential therapeutic use in treatment-resistant mood and anxiety disorders. Though it has been used by humans for centuries, research on PSI was curtailed for approximately 30 years following the US DEA’s assignment of psychedelic substances as Schedule I drugs in 1970, despite thousands of studies that showed promising results in the treatment of anxiety, obsessive-compulsive disorders, addiction, and sexual dysfunction (1). The advancement of magnetic resonance imaging (MRI) and positron emission tomography (PET) imaging in the 1990s improved our understanding of the mechanisms within the central nervous system, reigniting interest in psychedelics research, and leading to pilot studies and eventual FDA approvals for the therapeutic use of psychedelics in the treatment of various mental health conditions (2). Recent studies show efficacy in treating substance dependence(3), depression (1, 4), and anxiety(5).

Several clinical studies have been conducted with healthy human volunteers. de Veen et al., concluded that PSI may exert anti-addictive properties by relieving negative emotional states and stress or by lessening cognitive inflexibility and compulsivity(3). In a recent study, PSI has been suggested to have some use in the treatment of acquired brain injury by way of modulating excessive neuroinflammation and promoting neuroplasticity(6). It has also been shown to modulate self-focus, reduce negative emotional processing, and improve social functioning in healthy patients, as well as patients with psychiatric illness (1). Human studies are limited, however, in the range of experiments that can be performed using PSI, due to the lack of knowledge about the exact nature of the neurological mechanisms at play as well as ethical concerns about inducing a psychologically vulnerable state (6). This is an issue that has been partially addressed with research in mammals.

Using rodents, researchers have been able to identify the dose-dependent effects of PSI more precisely on the brain on a much shorter timescale. Jefson et al. showed the drug’s dose-dependent facilitation of plasticity-related gene expression in the rat prefrontal cortex and hippocampus(7). Similarly, it was shown to induce long-lasting increases in spine density and spine head width in the mouse medial frontal cortex as a result of a higher rate of dendritic formation (8). Increased neuroplasticity, or the structural and functional adaptation of neurons over time, has been hypothesized to alleviate symptoms of depression, a condition associated with the loss of synapses in the frontal cortex (8). This is one of the major mechanisms by which PSI is proposed to be of therapeutic use in the treatment of mental illnesses. Though a significant amount of information has been drawn from animal histology and behavioral studies, very few studies have performed MRIs on rodents; hence, there is a lack of a localized functional understanding of substances like PSI on the brain. Within that smaller amount of literature, there is an even smaller amount of imaging that has been done on awake as opposed to anesthetized rodents.

One such study illustrates the disruption of top-down processing as a result of PSI administration. Golden et al. (2022) found that PSI desynchronized activity in the anterior cingulate cortex (ACC) in mice by increasing the power of gamma oscillations (brain waves) and decreasing the power of low frequency oscillations(9). The resulting change in cortical activity and disruption of top-down processing is suggested to be the potential therapeutic mechanism of action of psychedelic compounds. In the present study we used BOLD imaging to record changes in brain function in response to PSI given to awake rats during the imaging session. We found dose-dependent global increases in positive BOLD volume of activation and hyperconnectivity with PSI treatment.

## Methods and Materials

### Animals

Adult male (n= 24) and female (n= 24) Sprague Dawley rats were purchased from Charles River Laboratories (Wilmington, MA, USA). Animals were housed in Plexiglas cages (two per cage) and maintained in ambient temperature (22–24°C). Animals were maintained on a reverse L-D cycle with lights off at 0900 hrs and studied during the dark phase when they are normally active. All experiments were conducted between 1000 and 1800 hrs to avoid the transitions between the L-D dark cycles. Food and water were provided *ad libitum*. All animals were acquired and cared for in accordance with the guidelines published in the NIH Guide for the Care and Use of Laboratory Animals. All methods and procedures described below were pre-approved by the Northeastern University Institutional Animal Care and Use Committee under protocol number 23-0407R. Northeastern University’s animal care and use program and housing facilities are fully accredited by AAALAC, International. The protocols used in this study followed the ARRIVE guidelines for reporting *in vivo* experiments in animal research (10). Animals were monitored daily over the duration of the study for general health, food and water consumption. A 15% loss in body weight was set as a humane endpoint.

### Psilocybin and ketanserin preparation and administration

Psilocybin was acquired through the National Institute on Drug Abuse (NIDA) and distributed by the Research Triangle Institute. On the day of imaging PSI was prepared in sterile saline (0.9% NaCl). To deliver drug remotely during the imaging session, a poly-ethylene tube (PE-20), approximately 30 cm in length, was positioned in the peritoneal cavity. The range of doses of PSI were taken from the literature(11–13). Based on body weight, rats were randomly assigned to one of four experimental groups: 1) vehicle, 2) 0.03 mg/kg, 3) 0.3 mg/kg and 4) 3.0 mg/kg of PSI. Each group consisted of 12 rats divided equally between males and females. Due to motion artifact five rats were excluded from the study. Two male rats from the vehicle treatment, one male from the low dose treatment and one male and one female from the high dose group. Following the original study with 48 rats, we ran a pilot study with 7 males Sprague Dawley rats to evaluate the effect of the 5HT2a receptor antagonist ketanserin on changes in brain activity in response to the 3.0 mg/kg dose of PSI. Rats were housed and acclimated as described above. One hour prior to imaging, rats were given an I.P injection of ketanserin (2.0 mg/kg in saline).

### Acclimation for awake imaging

To mitigate the stress associated with head restraint, rats underwent an acclimation protocol to familiarize them with the restraining system, which consisted of a head holder and body tube. This system was designed to include a cushioned head support, eliminating the need for ear bars and, in turn, reducing discomfort to the animals while minimizing any unintended motion artifacts. These acclimation sessions were conducted daily for five consecutive days. During these sessions, rats were briefly anesthetized with 1–2% isoflurane for placement into the restraining system. Their forepaws were fastened using surgical tape.

Once fully conscious, the rats were positioned within an opaque black box, a “mock scanner,” for 60 minutes. Inside the mock scanner, a tape recording of the MRI pulse sequence was played to simulate the environment of the magnet bore and the imaging protocol. Under these conditions there is a significant decreases in respiration, heart rate, motor activity, and plasma corticosterone levels when comparing the first and last acclimation sessions, as reported by King et al. ^16^. This reduction in autonomic and somatic signs of arousal and stress improved signal resolution and image quality.

### Image acquisition

Five to six rats were imaged in a day. Each day had a mix of the different experimental groups known by all the investigators. Rats were scanned at 300 MHz using a quadrature transmit/receive volume coil built into the rat head holder and restraining system for awake animal imaging (Ekam Imaging, Boston, MA. USA). A video of the rat preparation for imaging is available at www.youtube.com/watch?v=JQX1wgOV3K4. The design of the coil provided complete coverage of the brain from olfactory bulbs to brain stem. Radio frequency signals were sent and received with a quadrature volume coil built into the animal restrainer (Ekam Imaging, Boston MA, USA) ^(14)^. Imaging sessions were conducted using a Bruker Biospec 7.0 T/20-cm USR horizontal magnet (Bruker, Billerica, MA, USA) and a 2 T/m magnetic field gradient insert (ID=12 cm) capable of a 120-μs rise time. At the beginning of each imaging session, a high-resolution anatomical data set was collected using a rapid acquisition, relaxation enhancement pulse sequence (RARE factor 8); (25 slices; 1 mm; field of view (FOV) 3.0 cm^2^; data matrix 256 × 256; repetition time (TR) 3 s; echo time (TE) 12 ms; Effective TE 48 ms; number of excitations (NEX) 3; 4.48 min acquisition time; in-plane resolution 117.2 μm^2^). Functional images were captured using a multi-slice Half Fourier Acquisition Single Shot Turbo Spin Echo (HASTE) pulse sequence. This involved collecting 22 slices, each 1.1 mm thick, with the same FOV of 3.0 cm². The data matrix was 96 x 96, with a TR of 6 seconds, TE of 3.75 milliseconds, and an effective TE of 22.5 milliseconds. The acquisition took approximately 25 minutes, with an in-plane resolution of 312.5 μm². It should also be emphasized that high neuroanatomical fidelity and spatial resolution are critical in identifying distributed neural circuits in any animal imaging study. Many brain areas in a segmented rat atlas have in-plane boundaries of less than 400 μm^2^ and may extend for over 1000 μm in the rostral/caudal plane. With the development of a segmented, annotated 3D MRI atlas for rats (Ekam Solutions, Boston, MA, USA), it is now possible to localize functional imaging data to precise 3D “volumes of interest” in clearly delineated brain areas. This spatial resolution was sufficient to identify the bilateral habenula, with approximately 4–5 voxels on each side, but not to differentiate between the lateral and medial habenula.

### Data analysis

The fMRI data analysis consisted of three main steps: pre-processing, processing, and post-processing. All these steps were executed using SPM-12 (available at https://www.fil.ion.ucl.ac.uk/spm/) and in house Matlab software. In the pre-processing stage, several operations were performed, including co-registration, motion correction, smoothing, and detrending. Co-registration was carried out with specific parameters: Quality set at 0.97, Smoothing at 0.6 mm Full Width at Half Maximum (FWHM), and Separation at 0.4 mm. Additionally, Gaussian smoothing was applied with a FWHM of 0.8 mm.

The processing step involved aligning the data to a rat atlas, followed by segmentation and statistical analysis. To achieve registration and segmentation, all images were initially aligned and registered to a 3D Rat Brain Atlas©, which included 173 segmented and annotated brain regions. This alignment was performed using the GUI-based EVA software developed by Ekam Solutions (Boston, MA). The image registration process encompassed translation, rotation, and scaling adjustments, performed independently in all three dimensions. All spatial transformations applied were compiled into a matrix [Tj] for each subject. Each transformed anatomical pixel location was tagged with its corresponding brain area, resulting in fully segmented representations of individual subjects within the atlas.

Each scanning session consisted of 250 data acquisitions (NR) with a period of 6 sec (TR) each for a total lapse time of 25 min. The first 50 scans (5 min) were control window while the stimulation window was 200-240 (for ca 20 min post-injection) scans. Statistical t-tests were performed on each voxel (∼ 36,000 in number of voxels in the whole brain) of each subject within their original coordinate system. The baseline threshold was set at 1%. The t test statistics used a 95% confidence level (p<0.05), two-tailed distributions, and heteroscedastic variance assumptions. As a result of the multiple t-test analyses performed, a false-positive detection controlling mechanism was introduced. This subsequent filter guaranteed that the false-positive detection rate was below our cutoff of 0.05. The formulation of the filter satisfied the following expression:

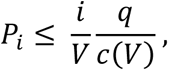

In the equation, Pi represents the p-value derived from the t-test conducted at the i-th pixel within the region of interest (ROI), comprising V pixels, with each pixel ranked according to its probability value. For our analysis, we set the false-positive filter value q at 0.2, and we fixed the predetermined constant c(V) at unity, following a conservative approach for assessing significance ^(15)^. Pixels that achieved statistical significance retained their relative percentage change values, while all other pixel values were set to zero. Our analysis employed a 95% confidence level, two-tailed distributions, and assumed heteroscedastic variance for the t-tests.

To create composite maps displaying the percent changes in the Blood Oxygen Level Dependent (BOLD) signal for each experimental group, we mapped each composite pixel location (in terms of rows, columns, and slices) to a voxel within the j-th subject using the inverse transformation matrix [Tj]-1. A trilinear interpolation method was used to determine the contribution of subject-specific voxel values to the composite representation. The use of inverse matrices ensured that the entire composite volume was populated with subject inputs. The average of all contributions was assigned as the percent change in the BOLD signal at each voxel within the composite representation of the brain for the respective experimental group.

In the post-processing phase, we compared the number of activated voxels in each of the 173 brain regions between the control and psilocybin doses using a Kruskal– Wallis test statistic. The data were ranked in order of significance, as detailed in **Tables 1 & 2**. We generated activation maps, depicted in **Fig 3**, showing brain areas with significant differences when comparing two or more groups.

### Resting State Functional Connectivity

#### Image Acquisition

Scans were collected using a spin-echo triple-shot EPI sequence (imaging parameters: matrix size = 96 x 96 x 20 (H x W x D), TR/TE=1000/15 msec, voxel size = 0.312 x 0.312 x 1.2mm, slice thickness = 1.2 mm, with 200 repetitions, time of acquisition 15 min. For preprocessing, we utilized a combination of various software tools, including Analysis of Functional NeuroImages (AFNI_17.1.12), the FMRIB Software Library (FSL, v5.0.9), Deformable Registration via Attribute Matching and Mutual-Saliency Weighting (DRAMMS 1.4.1), and MATLAB. Brain tissue masks for resting-state functional images were manually delineated using 3DSlicer and applied for skull-stripping. We identified motion outliers, which are data segments affected by substantial motion, and recorded the corresponding time points for later regression. Large motion spikes were also detected and removed from the time-course signals. Following this step, slice timing correction was applied to account for interleaved slice acquisition order. We performed head motion correction using the six motion parameters, with the first volume serving as the reference image. Normalization involved registering functional data to the 3D MRI Rat Brain Atlas © using affine registration through DRAMMS. After quality control, a band-pass filter (0.01 Hz to 0.1 Hz) was applied to reduce low-frequency drift effects and high-frequency physiological noise for each subject. The resulting images underwent detrending and spatial smoothing, with a full width at half maximum of 0.8 mm. Additionally, regressors, including motion outliers, the six motion parameters, the mean white matter, and cerebrospinal fluid time series, were incorporated into general linear models for nuisance regression to eliminate unwanted effects.

The region-to-region functional connectivity analysis was conducted to measure the correlations in spontaneous BOLD fluctuations. In this analysis, a network consists of nodes (brain regions of interest or ROIs) and edges (connections between regions). We averaged the voxel time series data within each node based on the residual images obtained through the nuisance regression procedure. Pearson’s correlation coefficients were computed across all pairs of nodes (14535 pairs) for each subject within all three groups to assess interregional temporal correlations. The resulting r-values, ranging from -1 to 1, were z-transformed using Fisher’s Z transform to improve their normality. We constructed 166 x 166 symmetric connectivity matrices, with each entry representing the strength of an edge. Group-level analysis was then conducted to examine functional connectivity in the experimental groups. The Z-score matrices obtained from one-group t-tests were clustered using the K-nearest neighbors clustering method to identify how nodes cluster together and form resting-state networks. A Z-score threshold of |Z| = 2.3 was applied to eliminate spurious or weak node connections for visualization purposes.

### 2.4.2 Functional Connectivity Analysis

#### Degree Centrality

We conducted all network analysis using Gephi, which is an open-source software for network analysis and visualization ^(16)^. We imported the absolute values of the symmetric connectivity matrices for both psilocybin and vehicle data, treating the edges as undirected networks. Degree centrality analysis measures the number of connections that a particular node has within the entire network. Degree centrality is defined as:

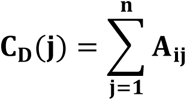

Here, “n” represents the total number of rows in the adjacency matrix denoted as “A,” and the individual elements of the matrix are indicated as “Aij,” which signifies the count of edges connecting nodes i and j.

### Statistics

We conducted all statistical analysis for the graph theory assessment using GraphPad Prism. To decide whether parametric or non-parametric assumptions were appropriate for different group subregions, we performed normality tests. We used Shapiro-Wilk’s tests to assess the normality assumption. Subregion degree centrality p-values exceeding 0.05 were considered to exhibit a normal distribution. Once the normality assumptions were confirmed, we employed one-way ANOVA to compare the degree centrality between the experimental groups in various subregions.

### Sex Differences

In an attempt to determine if there were sex differences in the sensitivity to PSI we divided the each of the experimental groups into their male and female cohorts. The final distributions were Vehicle (4 males, 6 females), 0.03 mg/kg (5 males, 6 females), 0.3 mg/kg (6 males, 6 females), and 3.0mg/kg (5 males, 5 females) PSI. While underpowered a two-way ANOVA comparing the sexes across all doses for whole brain activation was significant (p<0.0001) as reported in the Results. Dividing each experimental group into male and female cohorts for rsFC analysis was unsuccessful as the analysis requires a minimum of 7-8 subjects as determined from previous studies (17, 18)

### Psilocybin Assay

#### Sample Preparation

75 uL of plasma were added to 500 uL methanol (HPLC grade) in a centrifuge tube and spiked with 5 uL of 1 uM d4-psilocybin (Cayman). The 1 uM d4-psilocybin spike was prepared in a 20:80 methanol:water solution to match optimal elution conditions. Samples were briefly vortexed, after which they were incubated in the dark on ice for 30 minutes. After incubation, samples were centrifuged for 20 minutes at 19000 rcf at 20° C. Supernatant was added to 9.5 mL of HPLC grade water to form a 2.5% aqueous load. 1.5 mL of this was passed through a solid phase extraction (SPE) column (Agilent Bond Elut) and collected in a 2 mL autosampler vial. The remaining 8.5 mL of the load was passed through the SPE column and emptied to waste. 1 mL elutions were subsequently passed through the SPE column and collected in autosampler vials in the following concentrations: (water:methanol) 100:0, 80:20, 35:65: 25:75.

#### LC Properties

Mobile phase A consisted of 0.1% formic acid in 100% water (LCMS grade), and mobile phase B consisted of 0.1% formic acid in 100% methanol (LCMS grade). Mobile phase percentages were adjusted during the 10-minute run according adapted from Kolaczynska 2021(19). In brief, mobile phase A was started at 100% for 30 seconds. A linear ramp up to 100% B occurred from 0.5 minutes to 3 minutes. 100% B was maintained for one more minute (minutes 3-4), followed by a return to 100% A for the remaining six minutes.

#### Autosampler Conditions

The autosampler (Shimadzu SIL-40C) was kept at room temperature (25° C). Shimadzu pumps (SIL-40D) provided control of mobile phase percentages. The column oven, which housed the mixer and analytical column, was maintained at 40° C. A Phenomenex Luna C18(2) analytical column was used for chromatographic separation (50 mm length, 2 mm internal diameter, 3 uM particle size, 100 angstrom pore size). The analytical column was preceded by a Zorbax Eclipse XDB-C8 guard column.

#### Mass Spectrometric Analysis

Mass spectrometry was performed with a SCIEX 7500 triple quadrupole in positive Multiple reactions monitoring (MRM) mode with voltage set to 2300. Analysis of unknown peaks from plasma were quantified using standard curves as previously described(20) A mass spec method for psilocybin, d4-psilocybin, and psilocin was developed and parent/fragment pairs were optimized.

## Results

Shown in **Fig 1** are dose-dependent plasma levels of PSI and its active metabolite psilocin measure 30 min after IP administration. There was only four blood samples drawn for each dose and unfortunately the lowest dose of 0.03mg/kg was at the limit of sensitivity with only one sample providing a measure. The concentration in moles/liter is shown on a log scale to reflect the magnitude difference in each dose. The medium and high doses were significantly different (p<0.05). The 0.3 mg/kg dose gave a mean values of 0.07 µM and the 3.0 mg/kg dose 1.14 µM.

**Figure 1.**
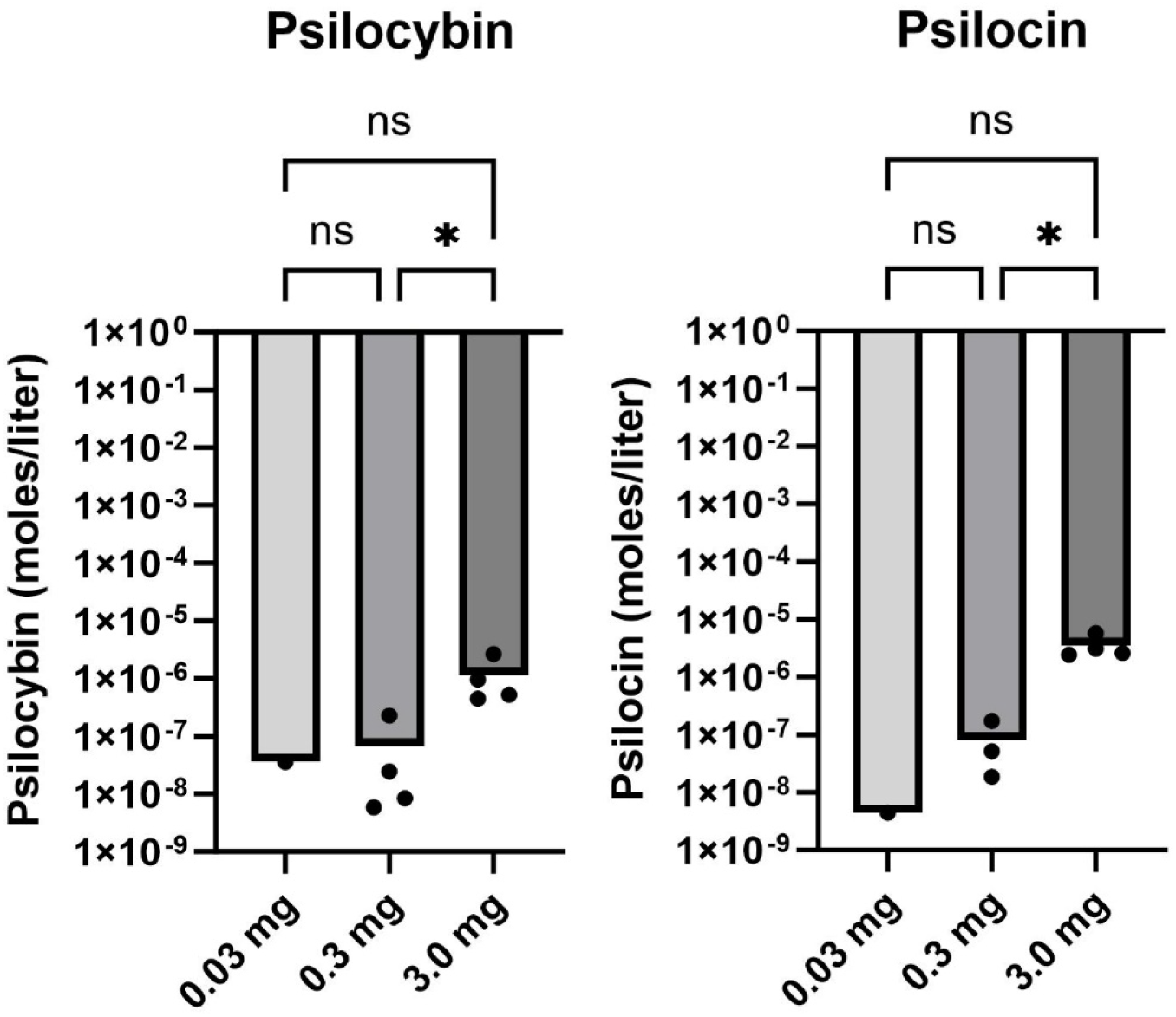
Plasma Levels of Psilocybin and Psilocin The dose-dependent plasma levels of PSI and its active metabolite psilocin are shown as bar graphs (mean) and dots (different samples) measured 30 min after IP administration. The concentration in moles/liter is shown on a log scale to reflect the magnitude difference in each dose. The medium and high doses were significantly different (p<0.05). The 0.3 mg/kg dose gave a mean values of 0.07 µM and the 3.0 mg/kg dose 1.14 µM.

After screening for acceptable motion artifact (see **Supplementary Fig 1**) rats were analyzed for volume of activation (VoA). See **Supplementary Excel File S1** for complete tables of VoA PSI dose response together with vehicle. **Table 1a**. is a truncated list of brain areas (mean ± SE) that show a significant dose-dependent change in positive VoA, i.e., number of positive BOLD voxels, in response to increasing doses of PSI. Significant positive activation was observed in 32/173 brain areas. The brain areas are ranked in order of their significance using a critical value of p<0.05. Shown are the p values and effect size given as omega square (Ω Sq). The false discovery rate (FDR) was p = 0.037. In **Table 1b**, only 7/173 areas displayed a significant change in negative VoA, an inconsequential effect when considering an FDR p = 0.008. Tables reporting the positive and negative VoA for all 173 brain areas for each dose of PSI are provided in **Supplementary Excel Files S2 & S3**.

**Table 1a.**
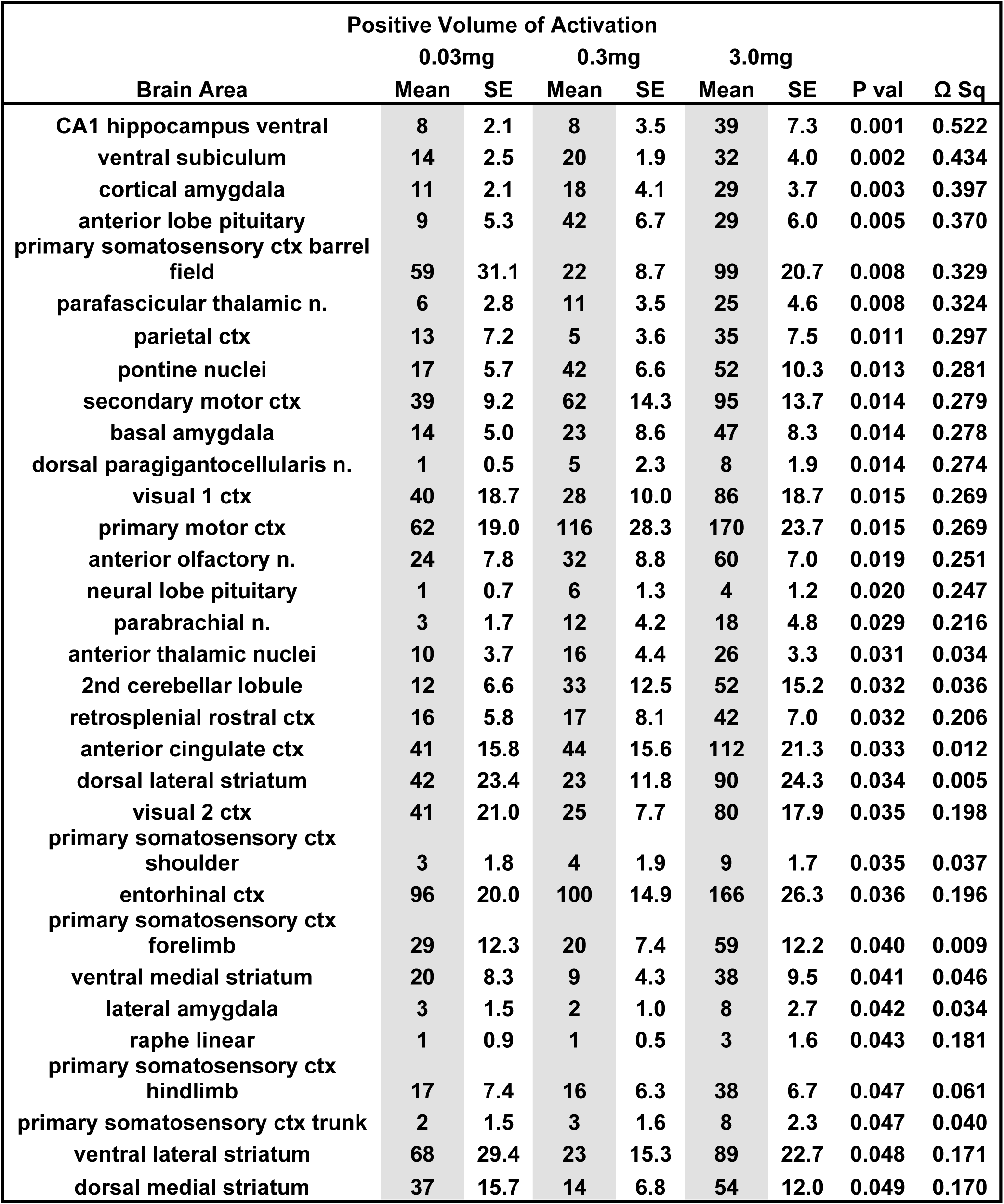
Psilocybin Dose Response: Positive Volume of Activation.

**Table 1b.**
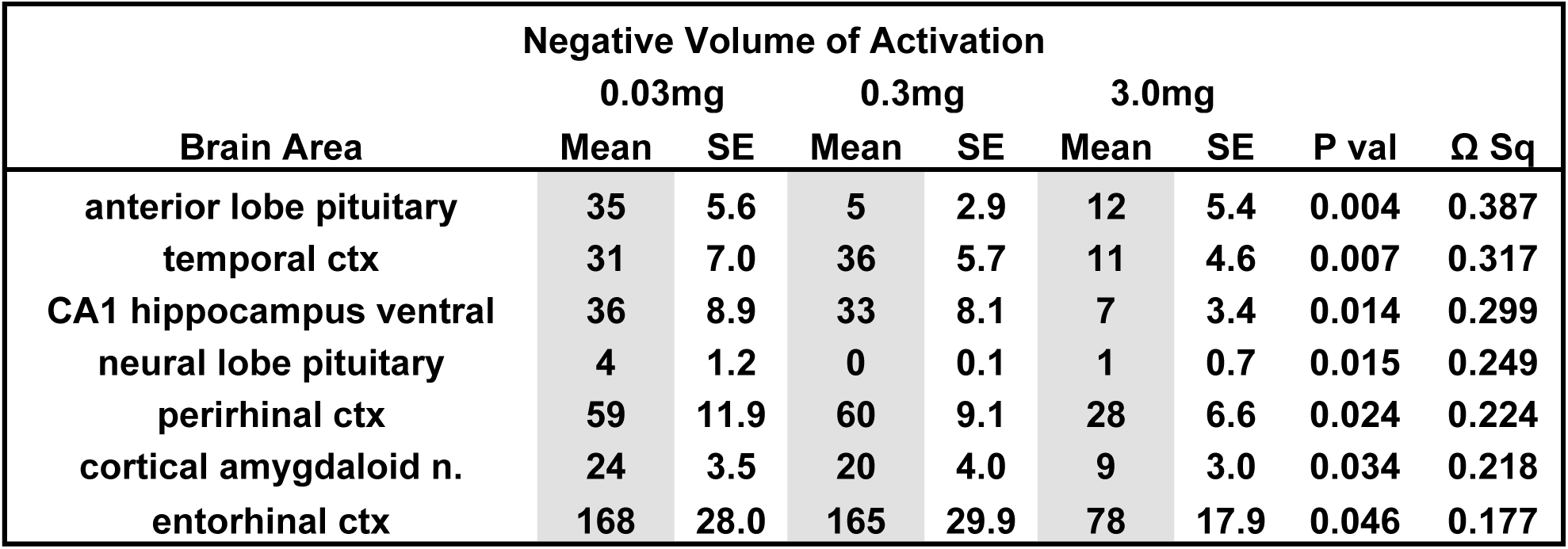
Psilocybin Dose Response: Negative Volume of Activation.

| Brain Area | Negative Volume of Activation |  |  |  |  |  |  |  |
| --- | --- | --- | --- | --- | --- | --- | --- | --- |
|  | 0.03mg |  | 0.3mg |  | 3.0mg |  | P val | Ω Sq |
|  | Mean | SE | Mean | SE | Mean | SE |  |  |
| anterior lobe pituitary | 35 | 5.6 | 5 | 2.9 | 12 | 5.4 | 0.004 | 0.387 |
| temporal ctx | 31 | 7.0 | 36 | 5.7 | 11 | 4.6 | 0.007 | 0.317 |
| CA1 hippocampus ventral | 36 | 8.9 | 33 | 8.1 | 7 | 3.4 | 0.014 | 0.299 |
| neural lobe pituitary | 4 | 1.2 | 0 | 0.1 | 1 | 0.7 | 0.015 | 0.249 |
| perirhinal ctx | 59 | 11.9 | 60 | 9.1 | 28 | 6.6 | 0.024 | 0.224 |
| cortical amygdaloid n. | 24 | 3.5 | 20 | 4.0 | 9 | 3.0 | 0.034 | 0.218 |
| entorhinal ctx | 168 | 28.0 | 165 | 29.9 | 78 | 17.9 | 0.046 | 0.177 |

A majority of brain areas (18/32) in **Table 1a** show a dose-dependent increase in VoA (gray highlight) e.g., ventral subiculum cortical amygdala, parafascicular n. Other areas like primary somatosensory ctx barrel field, parietal and visual cortices, dorsal lateral and ventral medial striatum show a U shaped dose response (11/32 areas) with the middle 0.3 mg/kg dose having the lowest VoA. These data suggest different brain regions e.g. basal ganglia, somatosensory ctx, prefrontal ctx, thalamus may show a different sensitivity to varying levels of PSI. Based on this observation all 173 brain areas were parsed into 11 different brain regions. The organization of these brain regions is shown in **Supplementary Excel File S4**. Shown in **Fig 2** are dot plots (individual brain areas) with bar graphs (mean ± SD) for the dose-dependent change in positive VoA for different brain regions. Note the U-shaped pattern of activation for the somatosensory cortex and basal ganglia where the medium dose of 0.3 mg/kg has the lowest effect. For the somatosensory cortex there was a significant treatment effect using a matched one-way ANOVA [F_(1.88, 35.80)_ = 110.7, p<0.0001)] followed by Tukey’s multiple comparison post hoc test * p<0.05; **** p<0.0001. There was a similar treatment effect and post hoc differences for the basal ganglia [F_(1.61, 11.28)_ = 149.7, p<0.0001), ***p<0.001, **** p<0.0001]. In all cases the high 3.0 mg/kg dose of PSI was significantly greater than the other doses for each brain region.

**Figure 2.**
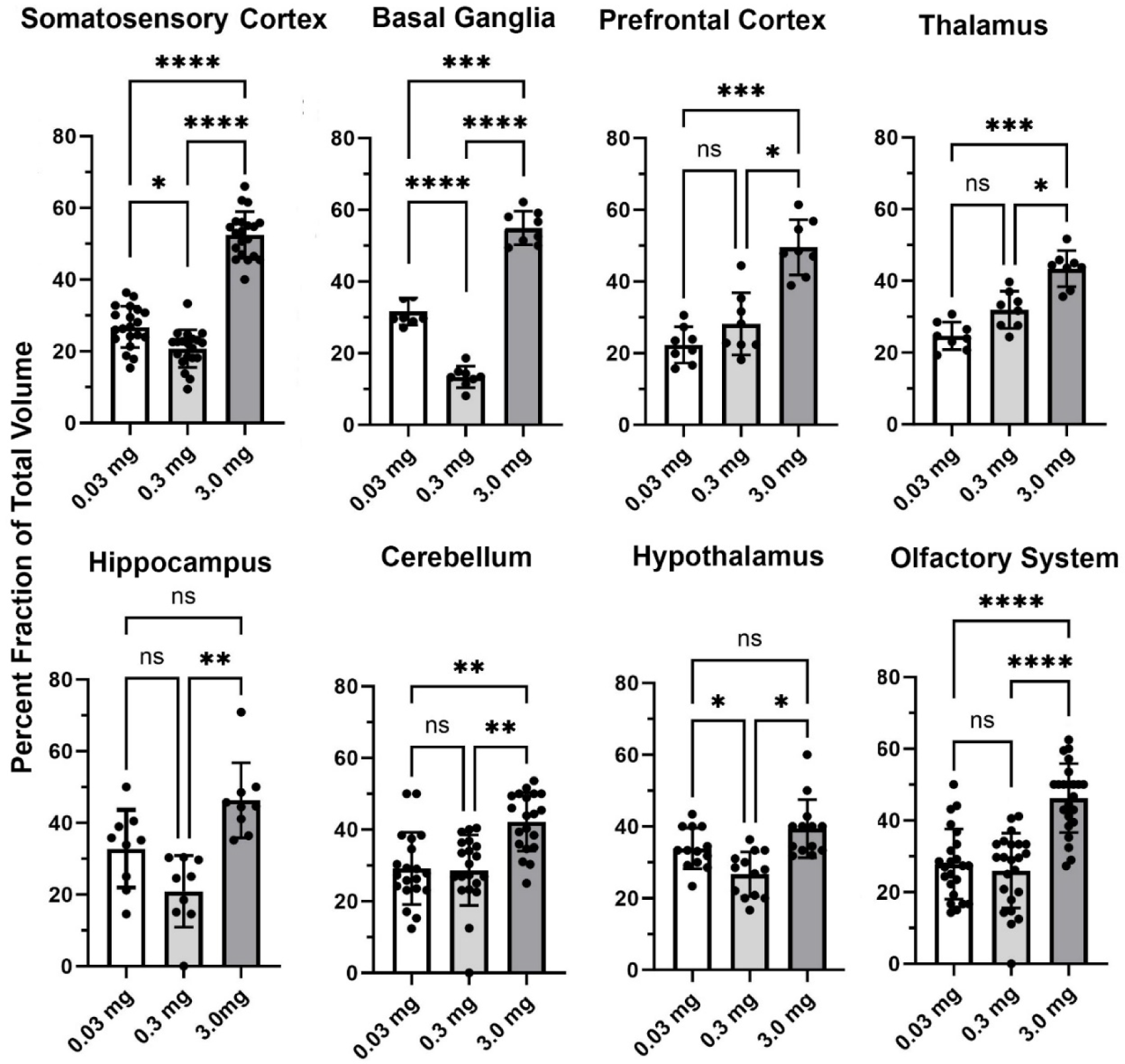
Regional Differences in BOLD Volume of Activation Shown are dot plots (individual brain areas) with bar graphs (mean ± SD) for the dose-dependent change in positive VoA for different brain regions. Data was compared using a matched one-way ANOVA followed by Tukey’s multiple comparison post hoc test. * p<0.05; ** p<0.01; ***p<0.001; **** p<0.0001.

Due to an outsized effect of the high dose, the 3.0 mg/kg group was used for further comparison with the vehicle group. **Tables 2a** and **2b** compare positive and negative VoA between the vehicle and high dose treatment, with the same reported metrics as **Table 1**. As such, results are also ordered by significance. An FDR analyses resulted in a p-value of 0.034. A total of 33 brain areas were significantly activated in the high dose group out of 173 brain areas observed. The pilot study pretreating rats with ketanserin and subsequently challenging them with the 3.0 mg/kg dose of PSI in the magnet showed a complete reduction in positive BOLD signal (**Supplementary Excel File S5**). Indeed, rather than an increase in BOLD signal there was a decrease in positive BOLD and an increase in negative BOLD to 13 different brain areas. When 5HT2a receptors are blocked the PSI effect is inhibitory to a small number of brain areas.

**Fig 3** shows the anatomical location of the brain areas listed in **Table 2a** for positive BOLD volume of activation presented as 2D statistical heat maps. The coronal sections are labeled (a.) through (k.) and arranged from rostral (top) to caudal (bottom). Areas in red are areas of significant activation following 3.0 mg/kg dose of PSI. Areas in yellow denote the location of white matter tracts. Brain section (a.) shows significant activity in the olfactory bulb. Sections (c.-d.) highlight afferent connections of the dopamine system (e.g., striatum, globus pallidus, anterior cingulate cortex), all producing significant activation. Sections (c.-f.) denote the somatosensory cortex, of significant interest due to its role in producing psychedelic hallucinations. Section (e.) highlights the thalamus thought to plays a role in psychedelic hallucinations. Note that sections (j.-k.) are not significantly affected by PSI. The essential findings taken from the 2D images, sections (a.- f.) are summarized in color-coded 3D reconstructions to the right. A transverse view of the brain shows the somatosensory cortex as a green translucent cover over all of the brain minus the olfactory bulb (yellow). A sagittal view of the brain displays the thalamus (blue) and basal ganglia (red).

**Figure 3.**
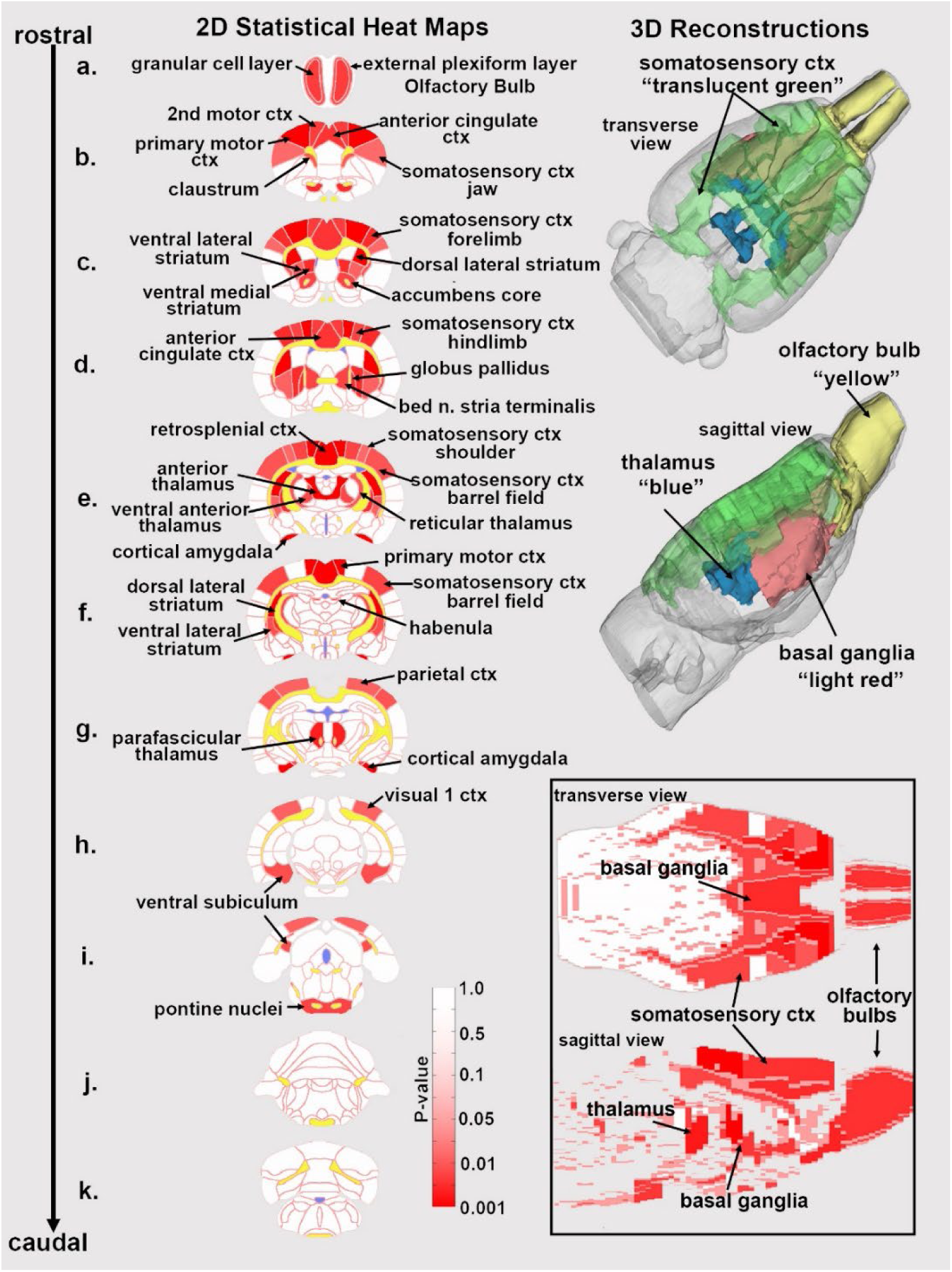
Statistical Heat Maps of Positive BOLD Activation Depicted are 2D coronal sections (a-k) showing the location of brain areas that were significantly different in positive BOLD VoA (red highlight) between vehicle and high dose (3.0 mg/kg) PSI. White matter tracts are highlighted in yellow. The insert in the lower right shows statistical heat maps in transverse and sagittal views. The 3D color-coded reconstructions summarize the major brain areas that were significantly different.

**Table 2.** Vehicle vs 3.0 mg/kg Psilocybin.

| Positive Volume of Activation |  |  |  |  |  |  |  |
| --- | --- | --- | --- | --- | --- | --- | --- |
| Brain Area | Vehicle |  |  | 3.0 mg |  | P val | Ω Sq |
|  | Mean | SE |  | Mean | SE |  |  |
| dorsal lateral striatum | 13 | 7.9 | < | 90 | 25.9 | 0.005 | 0.026 |
| retrosplenial rostral ctx | 10 | 3.5 | < | 42 | 7.5 | 0.005 | 0.476 |
| cortical amygdala | 12 | 3.5 | < | 29 | 3.9 | 0.007 | 0.435 |
| primary motor ctx | 76 | 13.6 | < | 170 | 25.3 | 0.007 | 0.017 |
| anterior thalamic n. | 10 | 3.8 | < | 26 | 3.6 | 0.008 | 0.011 |
| parafascicular thalamic n. | 8 | 3.3 | < | 25 | 4.9 | 0.009 | 0.037 |
| anterior olfactory n. | 27 | 7.1 | < | 60 | 7.4 | 0.013 | 0.356 |
| globus pallidus | 1 | 0.7 | < | 7 | 3.4 | 0.013 | 0.036 |
| primary somatosensory ctx forelimb | 23 | 7.4 | < | 59 | 13.0 | 0.015 | 0.041 |
| ventral medial striatum | 9 | 4.4 | < | 38 | 10.2 | 0.016 | 0.028 |
| anterior cingulate ctx | 36 | 12.5 | < | 112 | 22.8 | 0.017 | 0.320 |
| granular cell layer | 70 | 18.6 | < | 136 | 17.7 | 0.023 | 0.036 |
| secondary motor ctx | 49 | 10.3 | < | 95 | 14.6 | 0.023 | 0.285 |
| bed n. stria terminalis | 4 | 2.1 | < | 18 | 5.7 | 0.025 | 0.039 |
| pontine nuclei | 23 | 6.6 | < | 52 | 11.0 | 0.026 | 0.039 |
| primary somatosensory ctx hindlimb | 12 | 5.9 | < | 38 | 7.1 | 0.026 | 0.016 |
| accumbens core | 7 | 5.4 | < | 23 | 9.6 | 0.028 | 0.047 |
| ventral anterior thalamic n. | 2 | 1.2 | < | 6 | 1.4 | 0.028 | 0.043 |
| anterior pretectal n. | 4 | 1.1 | < | 12 | 2.7 | 0.030 | 0.254 |
| ventral subiculum | 19 | 5.0 | < | 32 | 4.2 | 0.034 | 0.238 |
| external plexiform layer | 31 | 6.2 | < | 55 | 8.5 | 0.039 | 0.046 |
| primary somatosensory ctx barrel field | 44 | 13.4 | < | 99 | 22.2 | 0.039 | 0.048 |
| white matter | 144 | 35.4 | < | 289 | 67.7 | 0.039 | 0.048 |
| claustrum | 1 | 0.7 | < | 5 | 1.5 | 0.041 | 0.047 |
| reticular n. thalamus | 6 | 2.2 | < | 15 | 3.3 | 0.043 | 0.046 |
| parietal ctx | 13 | 5.7 | < | 35 | 8.1 | 0.050 | 0.052 |
| ventral lateral striatum | 32 | 15.7 | < | 89 | 24.3 | 0.050 | 0.049 |
| visual 1 ctx | 44 | 13.1 | < | 86 | 20.0 | 0.050 | 0.048 |
| primary somatosensory ctx shoulder | 4 | 2.0 | < | 9 | 1.8 | 0.056 | 0.058 |
| primary somatosensory ctx jaw | 34 | 6.9 | < | 71 | 16.7 | 0.057 | 0.051 |
| dorsal medial striatum | 18 | 8.9 | < | 54 | 12.8 | 0.062 | 0.040 |
| primary somatosensory ctx trunk | 3 | 0.9 | < | 8 | 2.4 | 0.063 | 0.037 |

**Fig 4** shows the change in BOLD signal over time for the somatosensory cortex in response to vehicle (black line) and 3.0 mg/kg of PSI (red line). 250 images were acquired over the 25 min imaging session. Each acquisition is the mean ± SE of vehicle and high dose PSI rats combining the data from the somatosensory cortex (e.g. barrel filed, hindlimb, forelimb, trunk etc.) for all subjects for each experimental condition. With a two-way repeated measures ANOVA there was a significant interaction between time and treatment [F_(2.49, 202.42)_ = 4.424, p <0.0001] with a main effect for treatment [F_(1,117)_ = 10.55, p =0.0015] Note the immediate increase in BOLD signal (2%) following injection (image acquisition # 51) of PSI. There was a continuous increase in BOLD signal that remained above threshold (blue line) for almost the entire imaging session, peaking at 4% positive BOLD signal. In contrast, vehicle injection was characterized by a slow and steady increase in positive BOLD signal that did not breach the threshold until image acquisition #71, peaking at 2.5% positive BOLD signal. Shown in **Supplementary Fig 2** is the time course of BOLD signal change in the somatosensory cortices in response to 3.0 mg/kg PSI but in the presence of ketanserin. There is no significant difference between vehicle and PSI with the blockade of 5HT2 a receptors.

**Figure 4.**
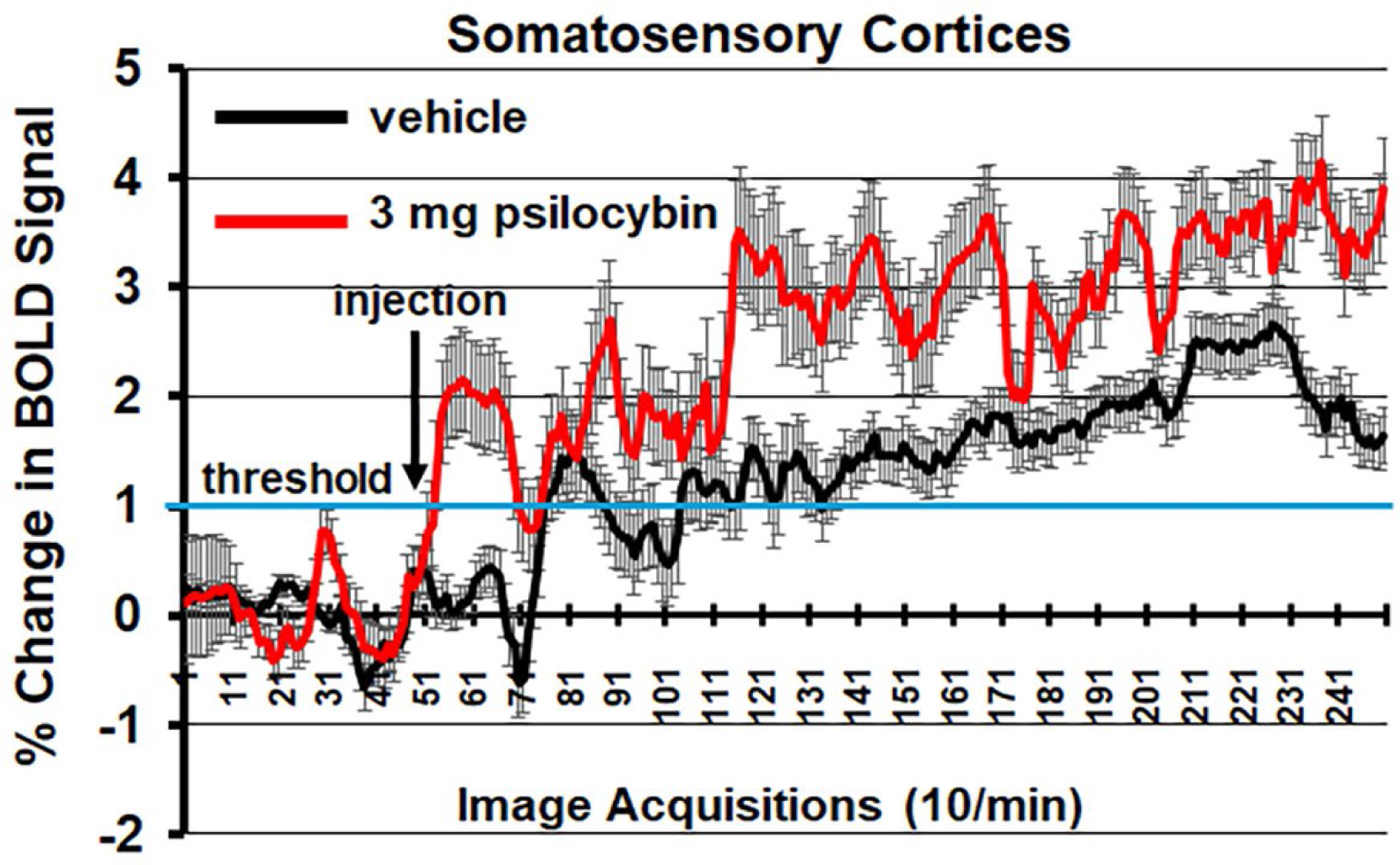
Somatosensory Cortex Time Course Shown is the change in BOLD signal over time for the somatosensory cortices in response to vehicle (black line) and 3.0 mg/kg PSI (red line). 250 images were acquired over the 25 min imaging session. Each acquisition is the mean ± SE of vehicle and high dose PSI rats combining the data from the different somatosensory cortices e.g. barrel field, hindlimb, forelimb, trunk, and upper lip. The 1% threshold is highlighted by the blue lines to account for the normal fluctuations in BOLD signal in the awake rat brain. A two-way ANOVA showed a significant Time x Treatment interaction F_(2.49, 202.42)_ = 4.424, p <0.0001.

When the mean volume of activation for each of the brain areas for male and female rats was compared for all experimental groups (veh, 0.03 mg/kg, 0.3 mg/kg, and 3.0 mg/kg PSI) there was a significant sex by dose interaction [two-way ANOVA F_(3,672)_ = 72.39, p<0.0001). Subsequent analysis (two-way ANOVA) looking at the sex difference in brain areas for each of the experimental groups showed no significant differences for vehicle [F_(169,1352)_ = 0,849, p=0.912)] or the 3.0 mg/kg dose of PSI [F_(170, 1360)_ = 0.9420, p= 0.686]. However, male were significantly more sensitive to the low 0.03 mg/kg dose of PSI as compared to females [mixed effects ANOVA, F_(169,1352)_ = 1.271, p = 0.0148]. A Wilcoxon signed-ranked test showed only six brain regions to be significantly activated. Females, on the other hand were far more sensitive to the 0.3 mg/kg dose of PSI as compared to males [two-way ANOVA, F_(169, 1690)_ = 3.273, p<0.0001]. A Wilcoxon post hoc test showed 45 brain areas were significantly activated. See **Supplementary Excel File S6** for a table of female/male differences for the 0.3 mg/kg dose of PSI.

Shown in **Fig 5** are the location of many of the brain areas in female rats that were more sensitive to the 0.3 mg/kg dose of PSI than males. Brain section (b.) shows a sex difference in the prefrontal cortex, e.g., prelimbic, ventral orbital and lateral orbital cortices. Section (c.) highlights the ventral striatum comprised of the ventral medial and ventral lateral striatum, accumbens core and shell, and ventral pallidum. Sections (d.- e.) note the sensitivity of the thalamus. Section (f.) shows the ventral tegmental area. Sections (f.- i.) highlight numerous areas comprising the ascending reticular activating system (e.g., midbrain reticular n., parabrachial n., paragigantocellularis and gigantocellularis reticular nuclei) together with the raphe nuclei and the periaqueductal gray, areas involved in arousal, sleep/wake cycles and consciousness. Sections (h.-i.) highlight a sex difference in the sensitivity of the cerebellum.

**Figure 5.**
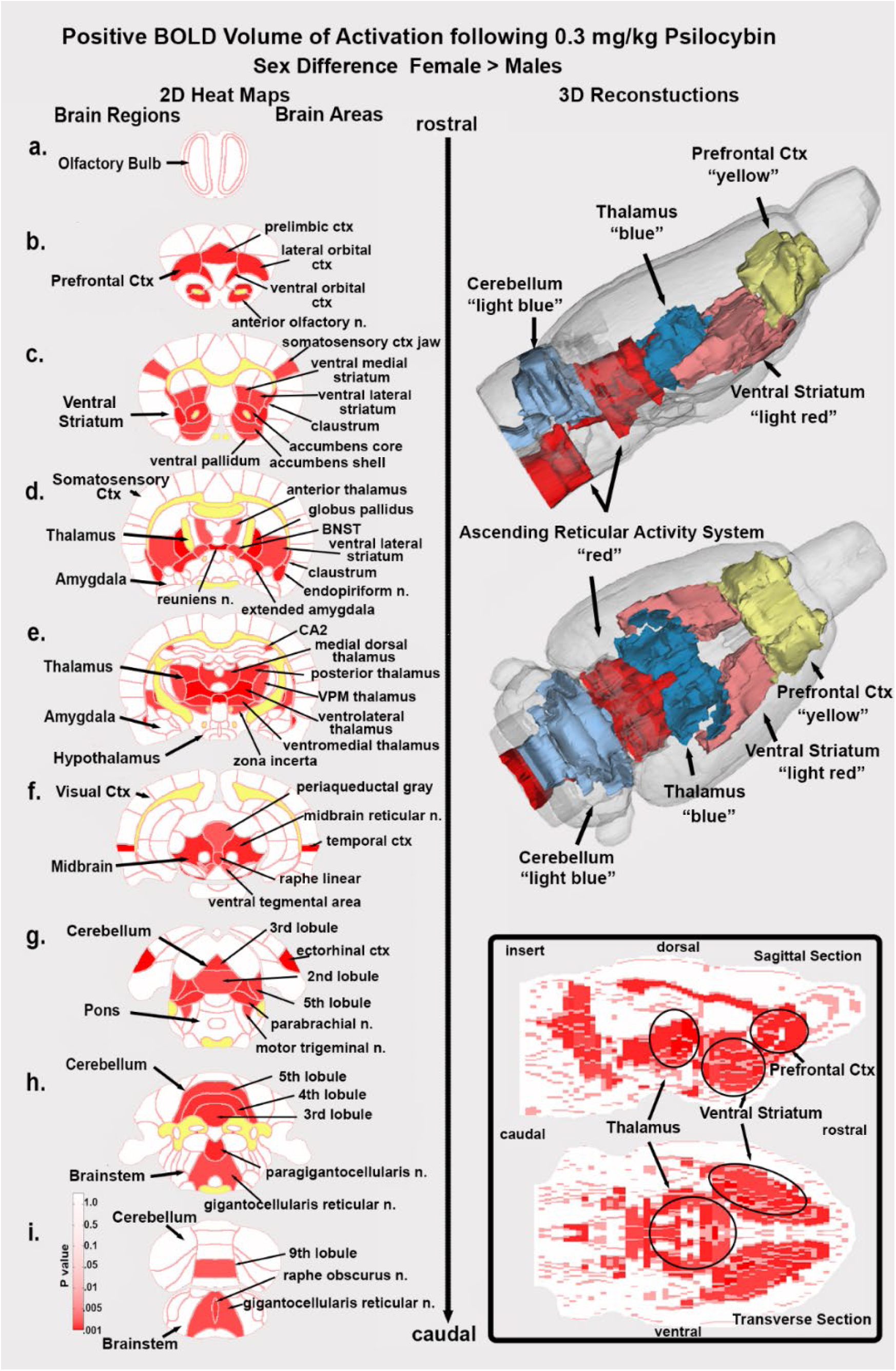
Sex Differences in Brain Activation Depicted are 2D coronal sections (a-i) showing the location of brain areas that were significantly different in positive BOLD VoA (red highlight) between males and females for the 0.3 mg/kg dose of PSI. Females were greater than males. White matter tracts are highlighted in yellow. The insert in the lower right shows statistical heat maps in transverse and sagittal views The 3D color-coded reconstructions summarize the major brain areas that were significantly different.

Shown in **Fig 6** are the dose-dependent differences in degrees (connections to other brain areas) for 11 major brain regions. The highlighted box in row A. shows the connections between all 173 brain regions as violin plots i.e., the probability distribution of the numerical data for each dose and vehicle (Veh, Avg Degree:10.082, Density:0.059; 0.03 mg/kg, Avg Degree: 22.971, Density: 0.135; 0.3 mg/kg, Avg Degree: 22.421, Density: 0.132; 3.0 mg/kg, Avg Degree: 25.474, Density: 0.150). There was a significant treatment effect using a matched one-way ANOVA [F_(2.60, 442.1)_ = 72.46, p<0.0001)] followed by Tukey’s multiple comparison post hoc test **p<0.01, *** p<0.001; **** p<0.0001. For many brain regions e.g. olfactory system, somatosensory cortex, amygdala, thalamus and midbrain all doses were significantly greater than vehicle, but not between themselves. In contrast in row B. the medulla [F_(2.76, 52.45)_=76.49,p<0.0001] and cerebellum [F_(2.76,52.52)_=72.46, p<0.0001] showed a significant difference between PSI doses and displayed a U-shaped profile with the medium 0.3 mg/kg dose having the lowest numbers of degrees. Note the high level of degrees (29.95 ± 10.1, mean and SD) for the low 0.03 mg/kg dose of PSI as compared to vehicle (6.05 ± 3.08). Row C. looks at these differences from the perspective of the medial cerebellar, dentate and interposed deep cerebellar nuclei. All efferent functional connections leaving the cerebellum and projecting to extra cerebellar sites go through these three nuclei. The significant difference (two-tailed t-test P=0.039) between vehicle and 0.03 mg/kg dose of PSI for the cerebellar nuclei is shown in the bar graph in row C. The individual connections between brain areas for vehicle and low dose PSI are shown as wheels with the three cerebellar nuclei in the middle. The cerebellar nuclei with vehicle treatment have only 15 connections between them show in red. With PSI treatment these connections are increase to 46 with the exception of the raphe linear and periaqueductal gray. The connections with PSI expand to include all of the cerebellum, areas of the hippocampus (dentate, subiculum, CA1), all of the olfactory bulb (glomerular, external plexiform and granular layers), temporal ctx, ventral medial thalamus, and dorsal medial striatum and many areas in the brainstem. The differences between vehicle and low dose PSI are shown in the color-coded 3D reconstruction below.

**Figure 6.**
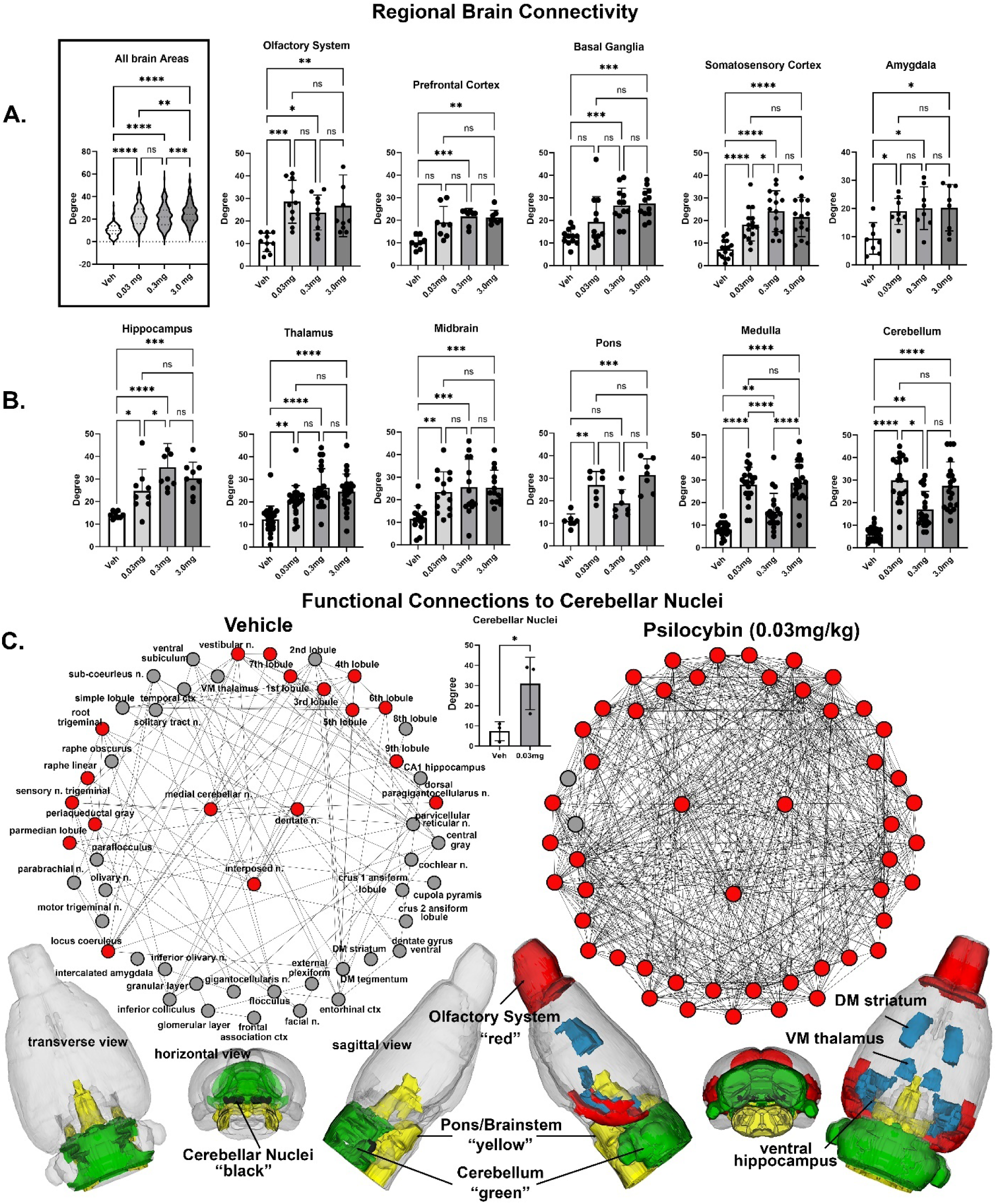
Regional Differences in Functional Connectivity Rows A. and B. shown are dot plots and bar graphs (mean ± SD) for the number of degrees or connections to other brain areas for veh 0.03 mg/kg, 0.3 mg/kg and 3.0 m/kg PSI. The violin plot show as an insert at the beginning of row A. is the total of all degrees between the 173 different brain areas. Shown in row C. re functional connections to the cerebellum. Shown are radial representations of the connections or degrees (lines) to the three cerebellar nuclei (center red), e.g., fastigial, dentate and interposed, following vehicle or 3.0 mg/kg PSI. The total network is represented by 52 brain areas as the union of both treatments. All brain areas highlighted in red have direct connections to the cerebellar nuclei under each experimental condition. The significant difference in connectivity (Student t-test, *p<0.05) is shown in the inserted bar graph at the center of the figure. The 39 brain areas comprising the network can be organized in three major brain regions - olfactory bulbs (red), cerebellum (green), and brainstem/pons (yellow), with connections to the thalamus, striatum and hippocampus (all in blue). The data are presented for different brain regions. *p<0.05; **p<0.01; ***p<0.001; ****p<0.0001

## Discussion

To the best of our knowledge this is the first pharmacological functional MRI (phMRI) study in animals or humans to show a dose-dependent change in BOLD signal in response to PSI. To make these data more relevant to the human condition, imaging was performed while male and female rats were fully awake without the confound of anesthesia and during the dark phase of L-D cycle when rats are normally active. There was a dose-dependent increase in BOLD signal in much of the forebrain, basal ganglia and somatosensory cortex and a global dose-dependent increase in functional connectivity. These findings are discussed with respect to imaging studies in healthy human volunteers exposed to PSI and their relevance, if any, to understanding the complex hallucinogen experience.

Psilocybin is in the chemical class of indoleamine hallucinogens that also includes DMT and LSD (2). PSI is relatively non-selective for serotonin receptors having a high affinity for 5-HT_1A_, 5-HT_2A_, and 5-HT_2C_ receptor subtypes (21, 22). The CNS effects of PSI are primarily attributed to activation of the 5-HT_2A_ receptor localized to pyramidal neurons in the somatosensory and prefrontal cortices(23). There is evidence that PSI activation of 5-HT_2A_ receptor in the cortex leads to a rise in extracellular levels of glutamate (24). PSI can also elevate levels of dopamine in the basal ganglia(25) and prefrontal cortex(12). The involvement of glutamate and dopamine signaling are two examples of down-stream effects of PSI that could contribute to the complex changes in brain activity and functional circuitry associated with the psychedelic experience. To that end, phMRI is agnostic and not wedded to any particular mechanism, providing a global map of site-specific changes in brain activity and connectivity.

### Acute BOLD Response

There was a dose-dependent increase in positive BOLD volume of activation in many brain areas. Areas comprising the different body-specific parts of the primary somatosensory cortex (e.g., barrel field, hindlimb, truck, forelimb) were activated. Indeed the primary somatosensory cortices showed both a significant increase in volume of activation as well as percent change in BOLD signal over time, two separate ways to assess functional activity. The only other animal phMRI study of which we are aware was conducted by Spain et al., on male rats give a low (0.03 mg/kg) or high (2.0 mg/kg) dose of psilocin during the scanning session (13). The studies were performed on anesthetized rats during the light phase of their L-D cycle when rats are at rest or sleeping. The low dose of PSI had little or no effect. However, the high dose of PSI caused a significant increase in BOLD signal in the olfactory system, limbic and visual cortices but a decrease in the somatosensory cortex, thalamus, and hippocampus. Using a laser scanning technique, these same rats were also studied for changes in localized blood flow to the barrel field of the somatosensory cortex in response to whisker stimulation in the absence and presence of PSI while also recording local field potentials from implanted electrodes in the same site. The 2.0 mg/kg dose of PSI enhanced blood flow during whisker stimulation, a response that was augmented by PSI. Paradoxically, the drug-evoked increase in blood flow occurred during a decrease in neural activity raising questions about neurovascular coupling in the presence of PSI. However, vascular reactivity in response to a hypercapnic challenge was normal when comparing vehicle and PSI treatments. Golden et al., reported recordings from the anterior cingulate cortex on awake mice showing an increase in neuronal firing in response to a dose of 2.0 mg/kg of PSI (9). In addition there was a decrease in low frequency band power but an increase in gamma band power evidence of cortical desynchronization. These studies by Spain and Golden showing regional increases in BOLD signal and electrical activity and ours showing a global increase to PSI are in sharp contrast to the initial human BOLD imaging study performed by Carhart-Harris and colleagues (26). In healthy volunteers with a history of hallucinogen exposure there was a sharp decrease in cerebral blood flow (CBF) measured with arterial spin labeling in all areas of the brain. The same study was repeated with BOLD imaging again finding a decrease particularly in the prefrontal cortex and anterior cingulate. The magnitude of the decrease in CBF and BOLD was inversely correlated with the subjective effects of PSI. In a subsequent publication on data collected from the same subjects, Carhart-Harris reported PSI enhanced BOLD signal in the visual cortex and somatosensory areas in response to memory recall enacted during the scanning session showing a difference between resting and task driven effects of PSI (27). However, the global decrease in hemodynamic measures of brain activity in humans under resting conditions is contrary to PET imaging studies using [F-18] FDG to follow glucose metabolism. Vollenweider and colleagues reported a global increase in glucose metabolism following PSI treatment with the greatest changes occurring in the prefrontal and anterior cortical areas(28). The increase in metabolic hyperfrontality was also reported by Gouzoulis-Mayfrank et al. in healthy volunteers in response to PSI exposure (29). The Vollenweider group followed up their PET imaging study with another using arterial spin labeling to follow changes in CBF in response to PSI (30). Correcting for global CBF in each subject, they reported regional differences in blood flow highlighted by an increase in the frontal cortex but a decrease in other brain areas. N,N-dimethyltryptamine (DMT) a hallucinogen with comparable chemistry and activity to PSI, when tested in human volunteers also increases CBF to the frontal, limbic, and anterior cingulate cortices (31).

### Sex Differences

Dividing each of the experimental groups into male and female cohorts enabled us to test for sex differences to the sensitivity of different doses of PSI. Neither the vehicle nor the high 3.0 mg/kg dose of PSI presented with a significant sex difference. However, the low 0.03 mg/kg dose of PSI revealed a modest (only six areas activated) but significant male sensitivity to this hallucinogen. More compelling was the sensitivity of females to the medium 0.3 mg/kg dose of PSI with 45 brain showing a sex difference. The anatomical organization of the female sensitivity was most interesting with clearly defined activation of the ventral tegmental area and its dopaminergic efferent connections to the ventral striatum, accumbens and ventral pallidum (see **Fig 5**). Almost all of the thalamus was similarly activated together with key areas of the ascending reticular activating system. What was missing from this pattern of global brain activation was almost the entire somatosensory cortex and pons and all of the hippocampus, amygdala and olfactory system. Behavioral studies showing a sex differences in response to PSI or psilocin are sparse. Tyls and coworkers treated male and female rats SQ with 0.25 mg/kg, 1.0 mg/kg or 4.0 mg/kg of psilocin and 15 min later measured locomotor activity in an open field (32). There was a dose-dependent decrease in locomotion for both males and diestrus females but not estrus females. However, when tested for acoustic startle for prepulse inhibition there were no sex differences. Adult male mice but not female mice treated IP with PSI in doses ranging from 0.1 mg/kg to 2.0 mg/kg show reduced voluntary ethanol consumption (33). In a recent study Effinger et al., reported the central amygdala of female rats is more sensitive to a 2.0 mg/kg dose of psilocin as compared to males in measures of c-fos immunostaining and photometry in response to an aversive stimulus (34). There is no clear evidence in the clinical literature of sex difference in sensitivity to PSI (35).

### BOLD Functional Connectivity

One of the more compelling findings in this study was the dose-dependent increase in global functional connectivity. Reinwald and colleagues in a cross-over experimental design, imaged females rats treated with 1.0 mg/kg PSI and vehicle during the MRI scanning session(36). The studies were done under anesthesia and during the light phase of the L-D cycle. Unfortunately, the cerebellum was not included in the study. There was an overall global decrease in connectivity with PSI treatment, in stark contrast to our findings. The most plausible explanation for this disparity is the confound of anesthesia and impact of circadian biology. The importance of circadian timing and brain function cannot be underestimated. Perivascular clearance of an MRI contrast agent from the brain of awake rats is regulated by the circadian clock being highest during the light phase and lowest during the dark phase of the L-D cycle (37). Brain temperature is a circadian rhythm entrained by the L-D cycle (38, 39). These concerns notwithstanding, the data from the Reinwald study was consistent with many human imaging studies that report a general decrease in intra-network connectivity, particularly in association areas like the default mode and salience networks in healthy volunteers treated with PSI(40–42). Preller et al., reported a time-dependent change in global connectivity in healthy volunteers following PSI treatment (43). At 20 min post treatment these was hyperconnectivity in the occipital cortex. At later time points, 40 and 70 min post treatment, the pattern of brain activity evolved into hypoconnectivity in association areas and hyperconnectivity in sensory networks. Preller and colleagues further demonstrated that the altered brain connectivity was positively correlated with brain areas high in 5-HT_2A_ gene expression and negatively correlated with the localization of 5-HT_1A_ gene expression. Preller’s findings align with ours, demonstrating that PSI increases global brain connectivity across sensory networks. In a just published study Siegel and coworkers followed healthy volunteers prior to, during, and three weeks following a high 25 mg/kg dose of PSI (44). PSI had a dramatic effect on disrupting rsFC particularly in the default mood network.

One brain area highlighted in this study and previously overlooked in both preclinical and clinical literature is the cerebellum. Its involvement was anticipated, given the numerous awake animal imaging studies demonstrating alterations in cerebellar activity following the administration of cannabinoids, psychostimulants, ketamine, opioids, and neuropeptides (45–49) and most recently the hallucinogen LSD (18). Traditionally, the cerebellum has been considered primarily responsible for motor coordination. However, it also plays a role in autonomic physiological functions such as heart rate, blood pressure, and respiration (50–53). Additionally, the cerebellum is recognized for its significant involvement in emotional and cognitive functions (54–56), feeding (57), and addiction (58). The cerebellum’s connectivity is unique, with all efferent information passing through the fastigial, interposed, and dentate nuclei. These nuclei have extensive bidirectional connections with various brain regions, including the thalamus, hypothalamus, limbic cortex, amygdala, hippocampus, and brainstem (59). The cerebellum also receives a substantial portion of its nerve connections from the vestibular complex, which relays auditory information from the ear to the cortex. In this study, cerebellar connections extended to the olfactory bulb, frontal association cortex, and hippocampus. This extensive network of connections and its involvement in various behavioral and sensory functions raise the question: could the cerebellum play a role in hallucinogenic effects that mimic psychosis? Notably, given the cerebellum’s involvement in cognitive, emotional, and sensory processes, it may contribute to the wide range of symptoms and cognitive impairments observed in schizophrenia (60).

## Limitations

This study would have benefited from a suite of behavioral assays measuring cognition, locomotion and emotionality for each dose of PSI. These measures would have been particularly relevant to understanding the region specific pattern of brain activation shown in females to the medium 0.3 mg/kg dose of PSI. The involvement of the ascending reticular activating system and dopaminergic neural circuitry would suggest an increase in motivation.

This study did not include pharmacokinetics. While we analyzed plasma for PSI and its metabolite psilocin these samples were collected at single time point 30 min following IP injection. The drug-induced changes in BOLD signal were recorded 20-24 min after IP injection while the rsFC data were collected ca 35 min post injection. Would the data have differed if we chose later time points e.g., 45-60 min post injection of PSI? In a recent studying looking at the pharmacokinetics of 1 and 3 mg/kg PSI in mice injected IP, the levels of plasma psilocin peak at 15 with a half-life of ca 30 min (61). Hence, waiting a longer period of time for the acute drug-induced effect of PSI may have been limiting. The rapid rise of psilocin in plasma may account for the early rise in the BOLD signal observed in the time course series shown in **Fig 5** for the somatosensory ctx.

## Data Interpretation

This phMRI study testing the effects of PSI on brain activity is not unlike many others used to characterize a drug’s effect on the brain (48, 62, 63), yet the robust dose-dependent increase in global BOLD signal and connectivity are unprecedented and at odds with much of the preclinical and clinical literature. Spain and coworkers raised concerns around the potential confounding effects of 5-HT agonists and vascular tone with respective to the hemodynamic changes that contribute to the BOLD signal. Psilocin, psilocybin’s active metabolite, primarily acts through the 5-HT_2A_ receptor. Psilocin also binds with lower, but potentially relevant, affinity for the 5-HT_1B_ and 5-HT_1D_ receptors. Serotonin receptors lining the vasculature may modulate vascular tone and causes subsequent changes to BOLD signal independent of metabolism. Spain and colleagues also noted that 5-HT_2A_ receptors on cortical inhibitory interneurons may induce vasoconstriction and vasodilation, constituting a scenario where increased neuronal activity may lead to either positive or negative BOLD. Conscious, unrestrained rats treated with the hallucinogen DOI (2,5-dimethoxy-4-iodoamphetamine), a 5-HT_2A_ receptor agonist, reduces blood flow to the cutaneous circulation of the tail through vasoconstriction, an effect blocked by ketanserin pretreatment (64). Whether this vasoconstriction in the peripheral cutaneous circulation generalizes to the cerebral vasculature is unknown. The dose-dependent increase in BOLD signal in this study is not consistent with a decrease in cerebral blood flow; however, it aligns with the decreases in cerebral blood flow and BOLD signal reported in the Carhart-Harris study (26).

How do we reconcile these differences? Imaging done in awake animals and during the circadian L-D cycle when they are normally active could account for the discrepancies in the acute BOLD response between our study and that of Spain and Reinwald. Drawing comparisons between awake rats and awake humans when trying to explain the complex psychedelic effect of psilocybin or any hallucinogen by the pattern of brain activity may be an unreasonable endeavor. Commercially bred rats are, by most measures, genetically identical. All share a common restricted environment minimizing cognitive, emotional and sensory experience. This neurobiological portrait of their environment is limited to a short window of life - only 3-4 months. The food they eat is identical. They are drug naïve. Hence the variance between experimental doses may be small enough to observe dose-dependent changes across subjects as shown here. All of these controlled conditions are ideal for preclinical discovery but impossible in the world of human neuroscience. The psychedelic experience, by all accounts, is a personalized, untethered, and unfiltered emergence of experience and perception leading to “oceanic boundlessness” and “ego dissolution”(1) This complex subjective experience cannot be modeled on a rodent. It is unlikely, the head twitch in a rat is a proxy for a hallucinogenic experience(65, 66). At face value it is a peripheral measure of a motor response to the CNS effect of 5-HT_2A_ receptor activation (67). Nonetheless PSI activation of 5-HT_2A_ receptor in this study did favor enhanced BOLD signal in the sensorimotor cortex and other brain regions with a high density of 5-HT_2A_ receptors. Indeed, ketanserin treatment in this study blocked the activation of the somatosensory cortices by PSI. Brain areas hypothesized to be involved in loss of sensory filtering and organization of sensory motor stimuli such as the claustrum(68) and the cortico-basal ganglia-thalamic-cortical loop are all affected by PSI in a dose-dependent manner(2). So much of the neuroanatomical circuitry described in the human literature is affected in this study but not in the direction or pattern reported in the clinic.

### Summary

To the best of our knowledge there has never been a published study - preclinical or clinical - showing a <u>dose-dependent</u> change in brain functional BOLD activity and resting state functional connectivity using PSI. Furthermore we show a dose and sex-dependent difference in BOLD activity. Females are more sensitive to PSI and present with a very region specific pattern of activation around the thalamus and dopaminergic neural circuitry. These data were registered a rat 3D MRI atlas with 173 different bilateral brain areas providing global site-specific changes in male and female rats. All imaging was done in fully awake rats without the confound of anesthesia and during the dark phase of the L-D cycle when rodents are normally active. These experimental conditions were done to mimic the experience of a human imaging study. The neural circuitry associated hallucinations in humans. e.g. the cortico-basal ganglia – thalamic - cortical pathway and the claustrum were activated in a dose-dependent manner. Moreover, the cerebellum showed a dose-dependent increase in hyperconnectivity to numerous brain areas.

## Acknowledgements

We thank the National Institute on Drug Abuse, Drug Supply Program and Research Triangle Institute for providing the psilocybin used in these studies. We also thank the Center for Translational NeuroImaging and the Dept Pharmaceutical Sciences for financial support.

---

**Supplementary Figure S1.**
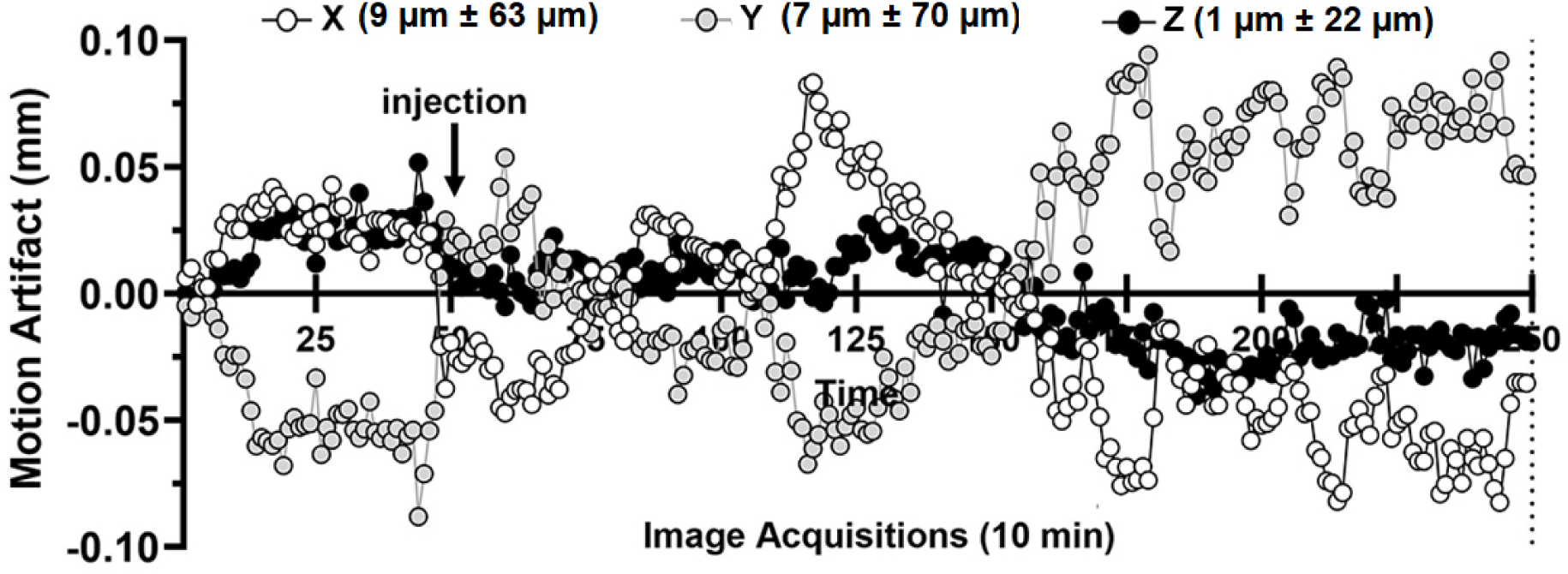
Motion Artifact Shown is a time course of motion artifact over the duration of the scanning protocol. The data from thirty-two rats (eight from each experimental group) were combined to show the mean ± SE for the X, Y and Z axis for 250 image acquisitions. The imaging parameters set the in-plane resolution of a pixel at was 312 µm^2^. The average motion artifact at any time point does not exceed 100 µm in any orthogonal direction.

**Supplementary Fig S2.**
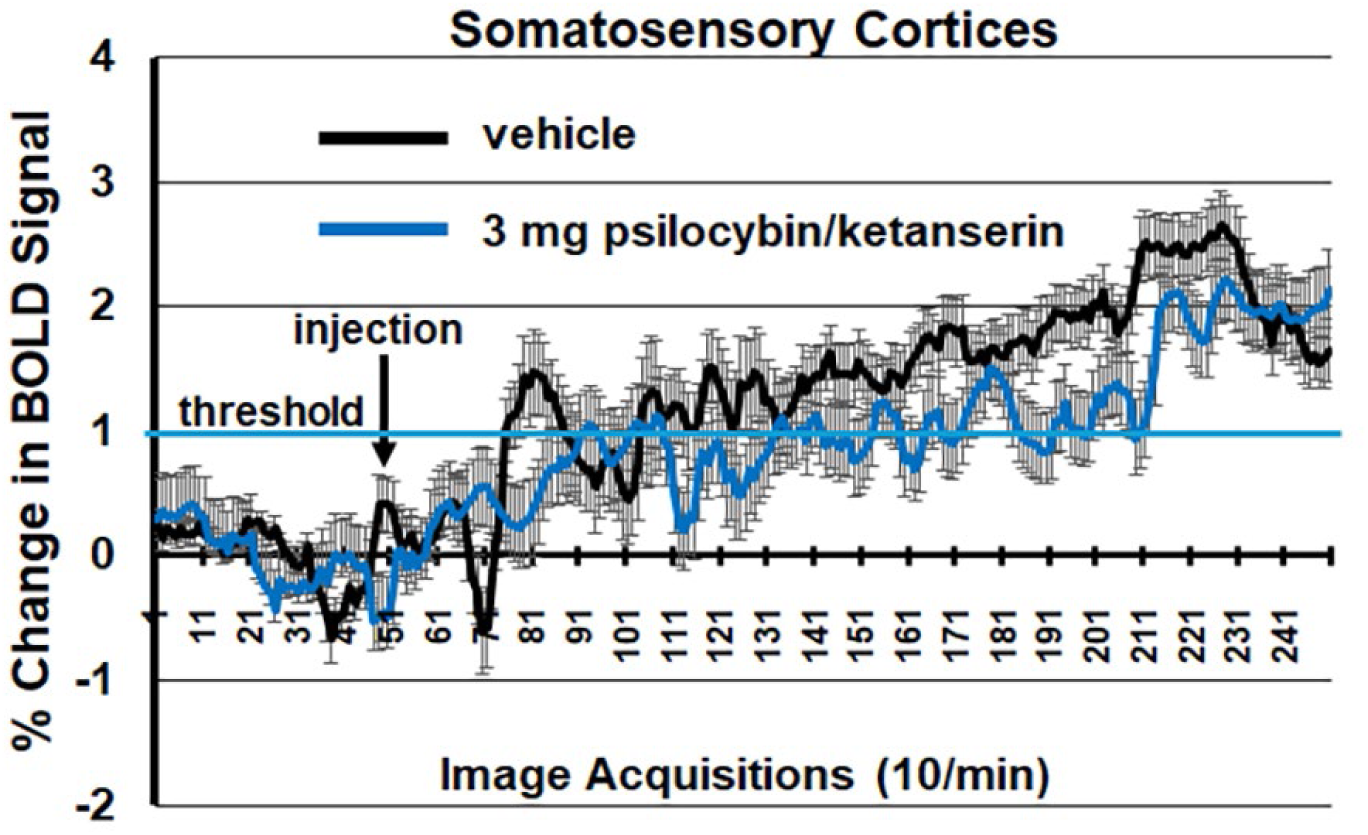
Shown is the time course of BOLD signal change in the somatosensory cortices in response to 3.0 mg/kg PSI but in the presence of ketanserin. There is no significant difference between vehicle and PSI with the blockade of 5HT2 a receptors.

